# Super-resolution single-cell spatial atlas of plant *de novo* regeneration

**DOI:** 10.64898/2026.02.20.705279

**Authors:** Xiehai Song, Shaoman Zhang, Zhiliang Yue, Yongqi Liu, Shanshan Chen, Yani Niu, Yan Shi, Hengjia Yang, Li Xu, Naixu Liu, Yuanyuan Miao, Man Lv, Jinshan Li, Tong Wang, Meizhi Xu, Binmei Sun, Chuan Qiu, Ruirui Xu, Jizong Wang, Huawei Zhang, Shuguo Hou, Gang Li, Haodong Chen, Xing Wang Deng, Bosheng Li

**Author notes:** These authors contributed equally (Xiehai Song, Shaoman Zhang, Zhiliang Yue, Yongqi Liu). Corresponding author: Bosheng Li Address: Peking University Institute of Advanced Agricultural Sciences, 699 Binhu Road, Weifang, Shandong, China.

## Abstract

Plant *de novo* regeneration, a capacity absent in animals, relies on stem-cell-niche formation rather than pre-existing reservoirs. Growth hormones such as auxin and cytokinin, together with numerous genetic regulators, participate in regeneration, but the architectural principles coordinating extensive cellular reprogramming at the individual-cell scale remain elusive. Using super-resolution multimodal spatial transcriptomics to 1.16 million cells, we tracked tomato regeneration across temporal intervals from wounding to organogenesis. Datasets are available at http://www.single-cell-spatial.com.

## INTRODUCTION

Regeneration, the capacity to replace damaged tissues or organs, serves as a fundamental pillar of biological resilience with striking evolutionary parallels across kingdoms [1, 2]. Though plants and animals use divergent cellular strategies, their regenerative processes appear governed by deeply conserved principles of cellular plasticity [3–5]. Animal regeneration frequently depends on pre-existing stem-cell niches (SCNs), yet dysregulation of these programs risks oncogenesis. Plants, conversely, show extraordinary regenerative potential by reprogramming somatic cells into *de novo* SCNs, capable of regenerating entire organisms while remaining tightly constrained to prevent aberrant growth [6–9]. This precise control establishes plants as powerful models for interrogating universal principles of cellular plasticity and tissue patterning.

Studies on plant regeneration have centered on callus formation, a pluripotent cell mass triggered by wounding or exogenous hormone application [10, 11]. Core roles are established for growth hormones (*e.g.*, auxin and cytokinin) and regulators such as *Wuschel* (*WUS*), *Regeneration Factor 1* (*REF1*), *Baby Boom* (*BBM*), the *Clavata* pathway (*CLV1/CLV3*), and *Somatic Embryogenesis Receptor Kinases* (*SERK1/2*) [4, 10, 12–16], but persistent cellular heterogeneity in callus has obscured our understanding of how spatial dynamics of positional cues and cell-fate patterns contribute to plant regeneration [17, 18]. Single-nucleus RNA sequencing (snRNA-seq), single-cell RNA sequencing (scRNA-seq) and spatial transcriptomics illuminate this complexity [19–24], however, compared with animal systems, plant models still lag in resolving SCN organization due to technical limitations in spatial resolution [25–30].

Here, we resolve this disparity through a super-resolution spatial atlas of plant regeneration, integrating 1.16 million single-cell profiles from 101 spatiotemporally resolved transcriptomic maps obtained following tomato wounding. As a resource, this atlas establishes a comprehensive, subcellular-resolution reference of plant regeneration and a standardized workflow, serving as a benchmark for spatial transcriptomics in plants, with potential relevance to other systems.

## RESULTS

### Multiscale spatiotemporal mapping of shoot regeneration after wounding

To investigate the spatiotemporal regulation of shoot regeneration, we conducted a multiscale analysis across tomato cotyledon explants from 0–16 days post-excision (DPE), a well-established *de novo* regeneration system used for genetic transformation, and noted for reproducible callus induction and characteristic morphology (Supplemental Figure 1) [10, 21]. Histological frozen sections (including transverse and longitudinal sections, Supplemental Figure 2A) captured distinct developmental phases: stem-cell establishment (0–8 DPE, with 0 DPE immediately after excision) and SCN formation (8–16 DPE), with spatial transcriptomics applied at 2 DPE intervals for most time points (Figure 1A–1C). We integrated sequencing-based spatial transcriptomics (using BGI stereo-seq V1, BMK S1000/S3000) to construct transcriptional landscapes [21], generating 60 spatial transcriptomes with a median detection of over 1,200 genes per cell at 5-μm resolution (Supplemental Figure 2B, Supplemental Data 1). Cell-type annotations were done by combining histological features with canonical marker-gene expression, validated by complementary snRNA-seq (10x Chromium, BMK DG1000) and bulk RNA-seq, thereby improving consistency among technologies and biological replicates (Figure 1C, Supplemental Figure 2B–2D, Supplemental Data 2).

**Figure 1.**
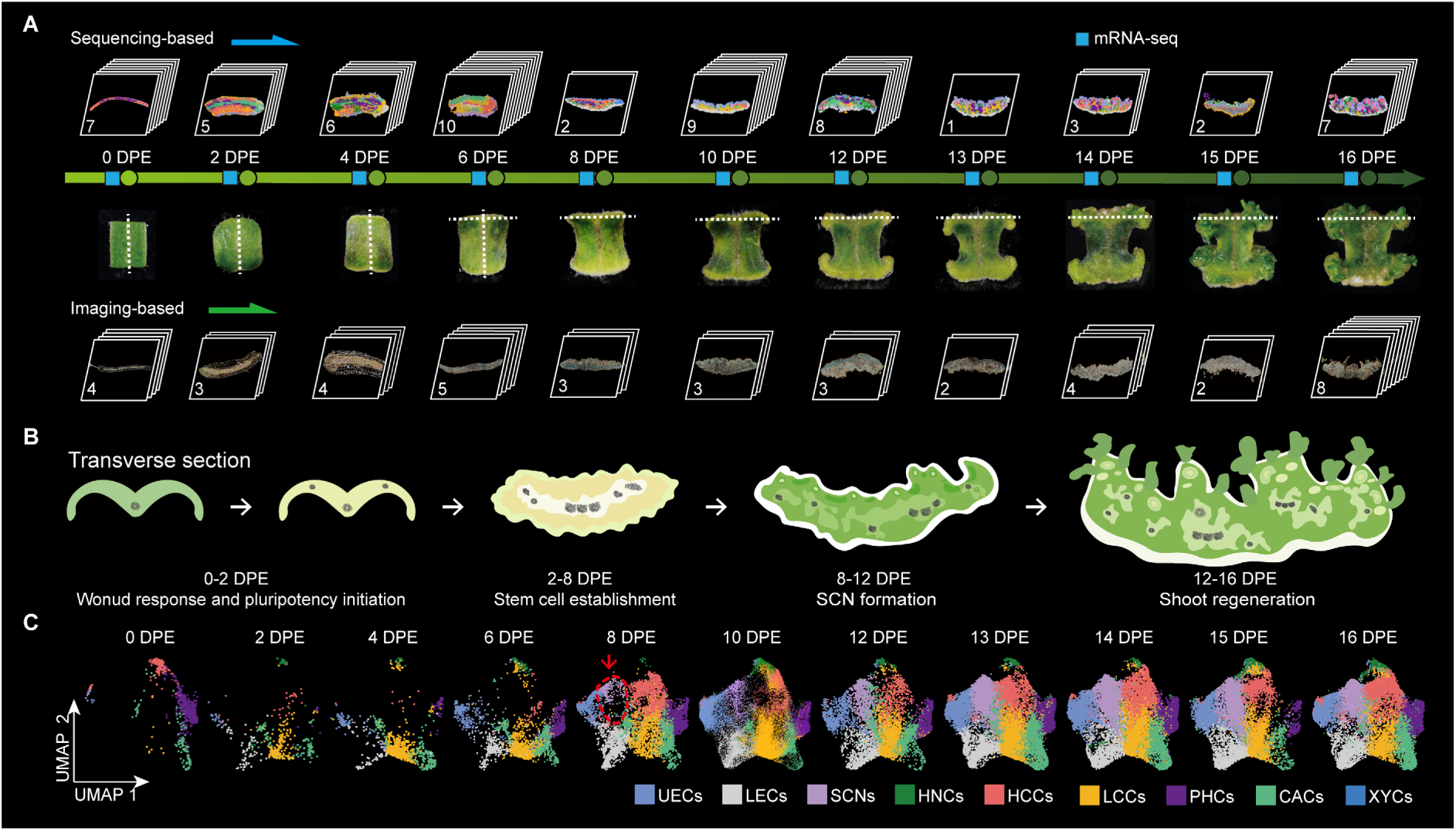
Construction of a spatial and cell-type atlas of plant regeneration. **(A)** Schematic workflow for constructing time-course spatial-transcriptomic profiles using sequencing-based, imaging-based, and bulk RNA-seq (square markers) approaches. Tomato explant (cotyledon) sections at developmental stages (0-16 DPE) are shown, with dashed white lines indicating sectioning planes. Numbered sections at each stage were used for transcriptomic profiling. **(B)** Model of callus development stages. Callus development is divided into sequential stages: 0 DPE (days post excision), explants immediately after excision; 0-2 DPE, wound response and initiation of pluripotency; 2-8 DPE, stem cell establishment; 8-12 DPE, stem-cell niche (SCN) formation; 12-16 DPE, shoot regeneration. **(C)** UMAP plots representing spatial transcriptomics of indicated time points after wounding. Cell-type classification: upper epidermal cells (UECs), lower epidermal cells (LECs), stem-cell niches (SCNs), hypoxic niche cells (HNCs), higher-chlorenchyma cells (HCCs), lower-chlorenchyma cells (LCCs), phloem cells (PHCs), cambium cells (CACs), and xylem cells (XYCs). The red arrows and dashed lines indicate the time periods when the SCN cell types appear.

**Figure 2.**
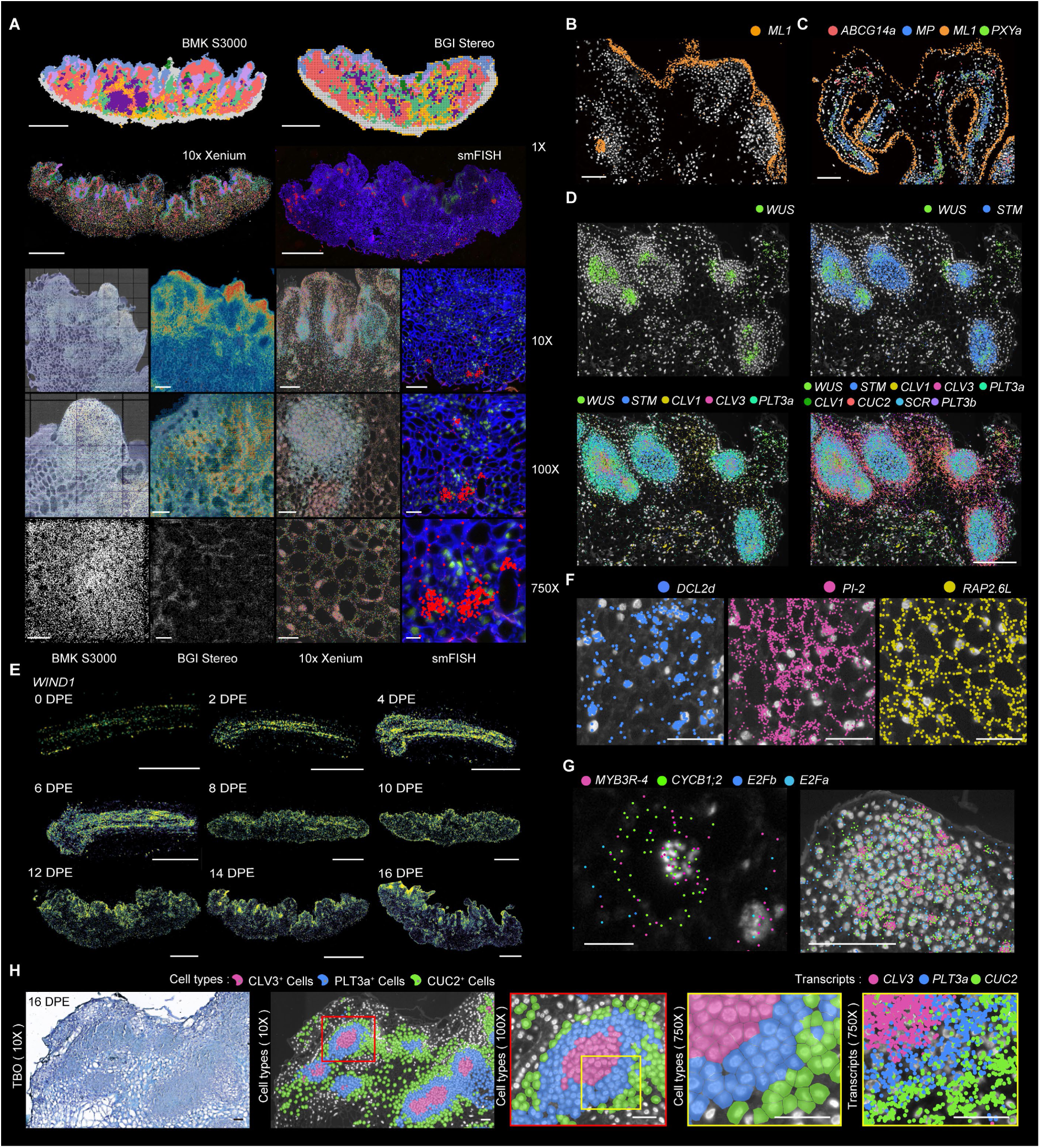
High-resolution spatial transcriptomics of tomato regeneration by 10x Xenium. **(A)** Multiscale spatial transcriptomics. At 1× magnification, cell clustering in callus tissues is shown using BMK S3000, BGI Stereo-seq (BGI Stereo), 10x Xenium, and smFISH (detecting *PROTEINASE INHIBITOR 2*, *PI-2* expression). Signal density maps of unique molecular identifier (UMI) counts for BMK S3000 and BGI Stereo, or imaging signal for 10x Xenium, are presented at 10×, 100×, and 750× magnifications. smFISH images reveal cell walls (blue), nuclei (green), and *PI-2* expression (red). Scale bars, 1× = 1 mm, 10× = 200 μm, 100× = 50 μm, 750× = 30 μm. **(B)** Visualization of *ML1* transcripts in the epidermis and SCN. Scale bar, 100 μm. **(C)** Spatial expression of cell-type markers detected by 10x Xenium. *ML1* defines the epidermis, while *ABCG14a*, *MP*, and *PXYa* mark vascular cells. Scale bar, 100 μm. **(D)** Co-detecting marker genes delineate SCN structure. Scale bar, 200 μm. **(E)** Temporal expression dynamics of *WIND1* from wounding (0 DPE) to complete shoot regeneration (16 DPE). Scale bar, 1 mm. **(F)** Spatial localization of *DCL2d*, *PI-2*, and *RAP2.6L* detected by 10x Xenium in 16-DPE tomato callus. Scale bar, 50 μm. **(G)** Left: Spatial visualization of cyclin gene expression detected by 10x Xenium. The G_2_/M phase marker genes *CYCB1;2* and *MYB3R-4* are enriched in the left cell, wherein chromatin condensation can be observed through nuclear fluorescence staining, indicating entry into the mitotic phase. Scale bar, 10 μm. Right: Single-cell resolution visualizes transcriptional dynamics within the shoot primordium. *E2Fa* and *E2Fb* mark gene the G_1_/S phase. Scale bar, 100 μm. **(H)** Subcellular-resolution profiling of tomato SCN structure using 10x Xenium at increasing magnifications. Markers: *CLV3* (core, cells in magenta, transcripts in pink), *PLT3a* (inner, cells in blue, transcripts in blue), *CUC2* (shell, cells in green, transcripts in green). Scale bar, 50 μm.

To address sparse gene detection and RNA spatial-coordinate drift, we incorporated 41 imaging-based spatial transcriptomics (10x Xenium) at subcellular resolution (∼30–200-nm probe spacing) [31] targeting 50 key genes identified from sequencing-based data related to hormone signaling, cell-type identity, developmental regulation, and cell-wall composition (Figure 1A, Supplemental Data 3). This expanded the total number of profiled cells in our atlas to 1.16 million. Overall, sequencing-derived gene lists informed probe selection for imaging tools, integrating genome-wide expression data with accurate cellular localization to construct a multimodal spatiotemporal atlas of plant regeneration. To facilitate community engagement, we developed the Plant Single Cell Spatial platform (http://www.single-cell-spatial.com), providing open access to datasets, thereby serving as a durable community resource for plant spatial transcriptomics (Supplemental Figure 3B–3G).

Our integrated spatial transcriptomics enabled mesoscale-resolution (resolution between subcellular- and tissue-level imaging) mapping of gene-expression during plant regeneration. At low magnification (1×), nine cell types were distinguishable, while higher magnification (10× and 100×) revealed localized expression aggregation in undifferentiated regions such as the SCNs. Subcellular imaging at 750× resolution visualized mRNA distribution within nuclear and peripheral cytoplasm compartments, with vacuolated regions exhibiting sparse signals (Figure 2A). At the single-cell level, spatial mapping highlighted dynamic cellular organization: *MERISTEM LAYER 1* (*ML1*) primarily marked epidermal cells but sporadically appeared in internal callus regions, reflecting irregular expression heterogeneity (Figure 2B) [32, 33]. Multi-gene co-expression patterns, including *ML1, PHLOEM INTERCALATED WITH XYLEM* (*PXY*) [34], *ATP-BINDING CASSETTE G14* (*ABCG14*) [35], and *MONOPTEROS* (*MP*/*ARF5*) [36] delineated differentiated cell types (Figure 2C). SCNs were further resolved using *WUSCHEL* (*WUS*) [13], *SHOOT MERISTEMLESS* (*STM*) [37], *PLETHORA 5* (*PLT5*) [38], *CUP-SHAPED COTYLEDON 2* (*CUC2*) [39], and *CLAVATA 3* (*CLV3*) [14], which defined functional subdomains (Figure 2D) [38, 40]. Time-course analysis captured gene-expression dynamics, such as *WOUND INDUCED DEDIFFERENTIATION 1* (*WIND1*) (Figure 2E) [12]. At the subcellular level, analysis uncovered compartment-specific gene activity during regeneration. *DICER-LIKE 2d* (*DCL2d*), involved in siRNA biogenesis [41], localized predominantly to nuclei, whereas *PROTEASE INHIBITOR 2* (*PI-2*) and *Related to APETALA 2.6-like* (*RAP2.6L*) [42] showed broadly distributed signals across both nuclear and cytoplasmic regions (Figure 2F). G_2_/M-phase cell-cycle regulators such as *CYCLIN B1;2* (*CYCB1;2*) [43] and *MYB DOMAIN PROTEIN 3R4* (*MYB3R-4*) [44] were enriched in nuclei with condensed chromosomes, contrasting with their sparse distribution in neighboring cells (Figure 2G) [45].

High-resolution analysis subdivided the SCN into distinct sublayers characterized by the expression of *CLV3*, *PLT3* and *CUC2* (Figure 2H, Supplemental Figure 3H). This comprehensive atlas, leveraging super-resolution spatial multi-omics, uncovering a greater degree of cellular heterogeneity whose function remains to be elucidated.

### Callus is generated from and connected to the original organ by vascular cells

Callus formation is known to depend on the ectopic activation of a lateral-root developmental program initiated by pericycle-like cells surrounding the vasculature [46–49]. Our atlas confirms this process and provides spatially resolved evidence with higher resolution and consistency than previously achieved, thereby validating established models in a tissue-wide context. At 0–2 DPE, a mitotically active zone marked by *CYCD3;1* and *CYCB1;2* expression emerged within vascular bundles and adjacent mesophyll cells (Supplemental Figure 4), representing the earliest proliferative domain. Meanwhile, the wound surface was sealed by a newly formed epidermis (Figure 3A). In parallel, *PLT*/*ANT* family genes (*PLT3a*, *PLT3b*, *PLT5*, *ANT*) were up-regulated in wound-adjacent regions, particularly near and continuous with pre-existing vascular bundles (Figure 3A). By 4 DPE, clusters of thin-walled callus cells appeared beneath the wound surface, consistent with a vascular cambium origin [46], coinciding with the expression of vascular regulators (*MP*, *PXYa*, and *VND7b*), indicating ongoing dedifferentiation (Figure 3A, Supplemental Figure 4 and 5A) [34, 36, 46, 50, 51]. Meanwhile, pluripotency-associated factors *PHB*, *REV*, *SHR* and *PLTs* remain highly expressed and become spatially stratified (Figure 3A and 3B, Supplemental Figure 4) [50]. Between 4–6 DPE, callus expansion continued, and reconnecting the severed vascular tissues, with transcriptional signatures indicative of new vascular element formation inside the callus. Newly formed callus cells adjacent to the cut surface continued to express vascular-associated markers *MP*/*PXY*, consistent with their association to pre-existing vascular tissues, as indicated by residual cut-site traces. Furthermore, *CUC2*, a key regulator of boundary formation and meristem organization, was expressed in the callus during this phase and subsequently showed increased expression as callus development progressed (Figure 3A and 3B).

**Figure 3.**
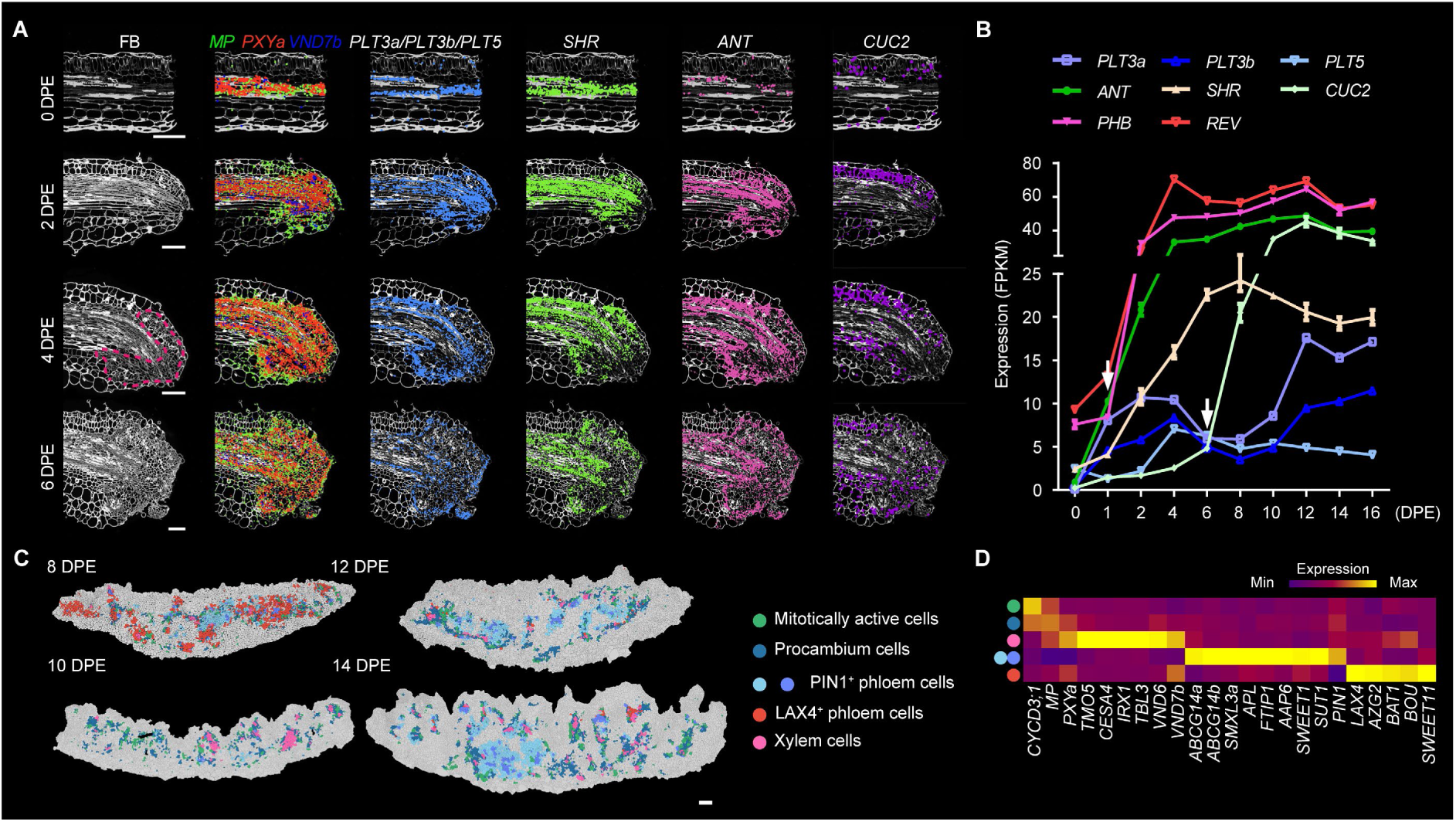
Developmental characteristics of vascular cells during regeneration. **(A)** Imaging-based spatial-transcriptomic mapping (10x Xenium) of vascular regulators (*MP*, *PXYa*, and *VND7b*) and pluripotency-associated factors (*PLT3a, PLT3b, PLT5*, *SHR*, *ANT*, and *CUC2*) transcripts from 2 to 6 DPE. Pink dashed circle represents callus initiation zone. Scale bar, 200 μm. **(B)** Bulk RNA-seq expression profiles of pluripotency-associated factors (*PLT3a, PLT3b, PLT5*, *ANT, CUC2*, *SHR*, *PHB* and *REV*) during regeneration (0-16 DPE). Arrows indicate induction of *ANT* at 1 DPE and *CUC2* at 6 DPE. Error bars indicate mean ± SD, n = 3 biological replicates. **(C)** Sequencing-based spatial transcriptomic (BMK S3000) maps showing spatial distribution of vascular sub-cell-types (procambium, mitotically active cells, xylem, PIN1^+^ phloem, LAX4^+^ phloem) across regeneration stages. Scale bar, 200 μm. **(D)** Bubble plots depicting the sequencing-based expression levels of select marker genes. The colors at left representing sub-cell-types are consistent with **C**.

The inner vascular cells exhibit transcriptional profiles consistent with specialized roles in transport and signaling during regeneration [21, 47]. Unsupervised clustering identified five vascular-cell populations: procambial cells, mitotically active cells, xylem cells, and two molecularly distinct phloem subtypes named PIN1^+^ and LAX4^+^ phloem cells (Figure 3C and 3D, Supplemental Figure 5B–5E, Supplemental Data 4). Each population has unique spatial distributions and transcriptional functional annotations: procambial cells are enriched in meristem maintenance and fate exchange; xylem cells execute lignin biosynthesis; and phloem subtypes diverge into roles for nutrient transport versus developmental signaling (Supplemental Figure 5F) [52, 53]. Transcriptomic profiling revealed phloem subtype specialization in hormonal and metabolic flux. The *PIN-FORMED 1* (PIN1^+^) phloem cells express auxin/cytokinin efflux transporters like *PIN1* [54], *ATP-binding cassette G14a/b* (*ABCG14a/b*) [55], and general amino-acid exporters like *Amino Acid Permease 6* (*AAP6*) and *Proton-dependent Oligopeptide Transporter 1* (*PROT1*), while *LIKE-AUX1 4*-enriched (LAX4^+^) phloem cells specialize in auxin/cytokinin influx transporters such as *LAX4 [56]* and *Azaguanine Resistance 2* (*AZG2*), as well as specific glutamate and alanine transporters like *BOUTON* (*BOU*) and *Bidirectional Amino Acid Transporter 1* (*BAT1*). Phloem subtype-specific sugar transporters, like *Sugars Will Eventually be Exported Transporter 1* (*SWEET1*) [57] and *Sucrose Transporter 1* (*SUT1*) [58], further indicating differential specialization in metabolite transport and allocation (Figure 3C, Supplemental Figure 5C–5E). The procambium, often considered a vascular stem-cell reservoir, displayed transcriptional profiles distinct from canonical SCNs (Supplemental Data 5). Procambium cells express stemness genes involved in vascular development, including *WUSCHEL-related Homeobox 4* (*WOX4*) and *CLAVATA3/embryo surrounding region-related 3* (*CLE3*) (Supplemental Figure 5G and 5H) [59]. Together, these findings highlight vascular cell diversity and suggest functional compartmentalization that may coordinate nutrient allocation, hormonal signaling, and aspects of developmental regulation during plant regeneration.

### A sub-cell-type of the upper-epidermal cells is pluripotent

During regeneration, the outer epidermal cells exhibit pronounced spatial heterogeneity, with unsupervised clustering distinguishing upper (air-exposed) and lower (medium-contacting) epidermal cells (Figure 1C, Supplemental Figure 2B). This distinction reflects tissue orientation in culture and is different from the dorsoventral polarity of leaves [60, 61]. Upper epidermal cells display a closer spatial association with SCNs (Figure 4A) and were further stratified into four subpopulations (Figure 4B-4E, Supplemental Data 6). Besides canonical epidermal cells related to cutin and wax, cells expressing *WUS*, a homeodomain transcription factor essential for plant meristem function [13], were classified as WUS^+^ cells, which were relatively abundant during early developmental stages but gradually declined over time (Figure 4B and 4D). Spatial transcriptomic analysis showed that *WUS* expression co-localized with the epidermal markers *ML1* and *PDF1* at early developmental stages, whereas at later stages their expression domains became spatially separated but remained adjacent (Figure 4F and G). This dynamic spatial relationship suggests that ML1 and PDF1 may regulate *WUS* expression.

**Figure 4.**
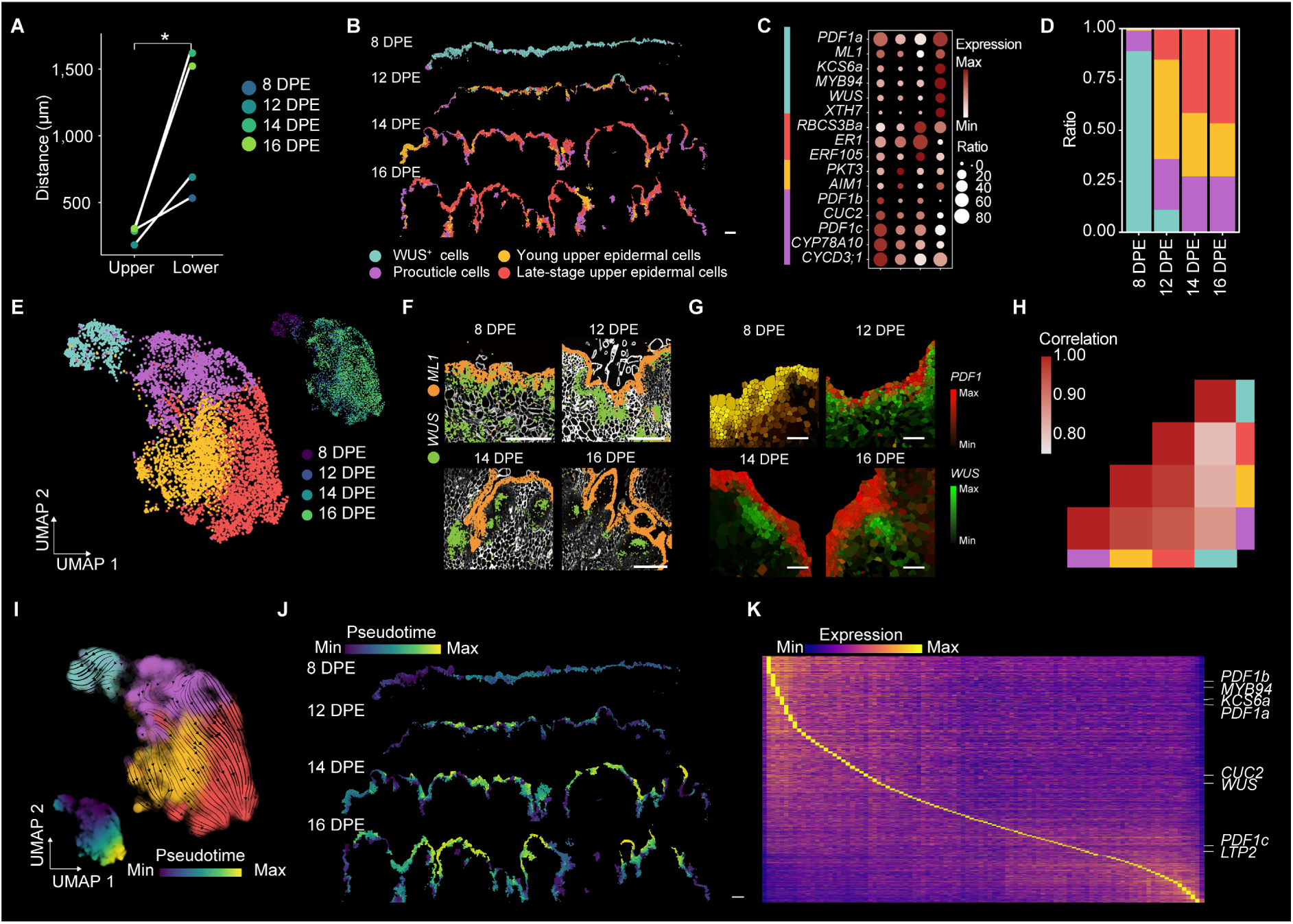
Developmental and functional heterogeneity of upper epidermal cells. **(A)** Distances between SCN cells and upper (air-exposed) versus lower (medium-contacting) epidermal cells at 8-16 DPE, quantified from sequencing-based spatial transcriptomics (BMK S3000). *, *P* < 0.05 (two-tailed *t*-test). **(B)** Spatial mapping of upper epidermal sub-cell-types, showing WUS⁺ pluripotency-associated cells, procuticle cells, young upper epidermal cells, and late-stage upper epidermal cells. Scale bar, 200 μm. **(C)** Bubble plot of marker-gene expression specific to sub-cell-types of upper epidermal cells. Sub-cell-type colors are consistent with **B**. **(D)** Proportions of the four sub-cell-types across 8-16 DPE, based on sequencing-based spatial transcriptomics (BMK S3000). Box colors representing sub-cell-types are same as in **B**. **(E)** UMAP plots of upper epidermal sub-cell-types colored by cell identity (left) or sampling time (right). Sub-cell-type colors are consistent with **B**. **(F)** Imaging-based spatial-transcriptomic (10x Xenium) mapping of *ML1* (orange) and *WUS* (green) transcripts from 8 to 16 DPE. Scale bar, 200 μm. **(G)** Sequencing-based spatial-transcriptomic (BMK S3000) mapping of *PDF1* and *WUS* expression from 8 to 16 DPE. Scale bar, 100 μm. **(H)** Heatmap of spatial gene-expression-profile correlation (Pearson correlation coefficient) between upper epidermal sub-cell-types and SCNs. Box colors are as in **B**. **(I)** RNA-velocity streamline plots (upper right), colors consistent with **E** and pseudotime (lower left) predicting the trajectory of upper epidermal cell lineage transitions. Colors are as in **B**. **(J)** Spatial projection of pseudotime trajectories for upper epidermal cells, corresponding to their predicted trajectories shown in **I**. Scale bar, 200 μm. **(K)** Heatmap of pseudo-temporal transitions of gene expression involved in upper epidermal cell development. Selected marker genes are annotated.

Transcript profiles of WUS^+^ cells diverged markedly from other epidermal subtypes (Figure 4H). RNA velocity [62] and Xenium-based spatial mapping supported the segregation of WUS^+^ cells from other epidermal populations, suggesting a role in SCN initiation and highlighting their distinct trajectory (Figure 4I and 4J). However, we note that some WUS⁺ cells may have arisen in a patchy and independent manner within the pluripotent cells rather than through relocation [1–3]. Pseudo-time analysis [63] revealed a transcriptional shift from pluripotency-associated genes (*WUS*, *CUC2*) [38, 39] to genes associated with differentiation and specialized cellular functions, consistent with a transition from pluripotency toward differentiation and underscoring dynamic transcriptional reprogramming linked to spatial positioning (Figure 4K). These results indicate that the upper epidermal cells prioritize defense and structural integrity and can also acquire pluripotency associated transcriptional states and stem-cell-like characteristics [64, 65].

### WUS^+^ cells in the pluripotent cells develop into the SCN core

Spatial-transcriptomic profiling classified four distinct populations within the pluripotent cells: SCNs, hypoxic niche cells, and two photosynthetic chlorenchyma subtypes of high- and low-photosynthetic gene-enriched cells (Figure 1C and 5A, Supplemental Figure 6A). Chlorenchyma cells predominate throughout the regeneration process, performing light-driven metabolic functions, while the sparse hypoxic niche cells are proposed to facilitate adaptation to low oxygen and oxidative stress [21, 66]. SCNs emerge and expand during 8–16 DPE and showed transcriptomic enrichment of proliferation-associated pathways (Figure 1C, Supplemental Figure 6B) [23], and comprise specialized subpopulations, notably a WUS^+^ cluster with transcriptional features consistent with niche progenitor identity (Figure 5B–5E, Supplemental Data 7) [4, 67, 68]. These cells share similar *WUS* expression with upper-epidermal WUS^+^ cells but differ in developmental regulators: SCN WUS^+^ cells up-regulate *CUC2* [39], associated with niche structure, whereas upper-epidermal WUS^+^ cells express the epidermal-differentiation marker *ML1* and *PDF1* [32, 33], suggesting that spatial context may influence fate specification (Figure 5F and 5G, Supplemental Data 8).

**Figure 5.**
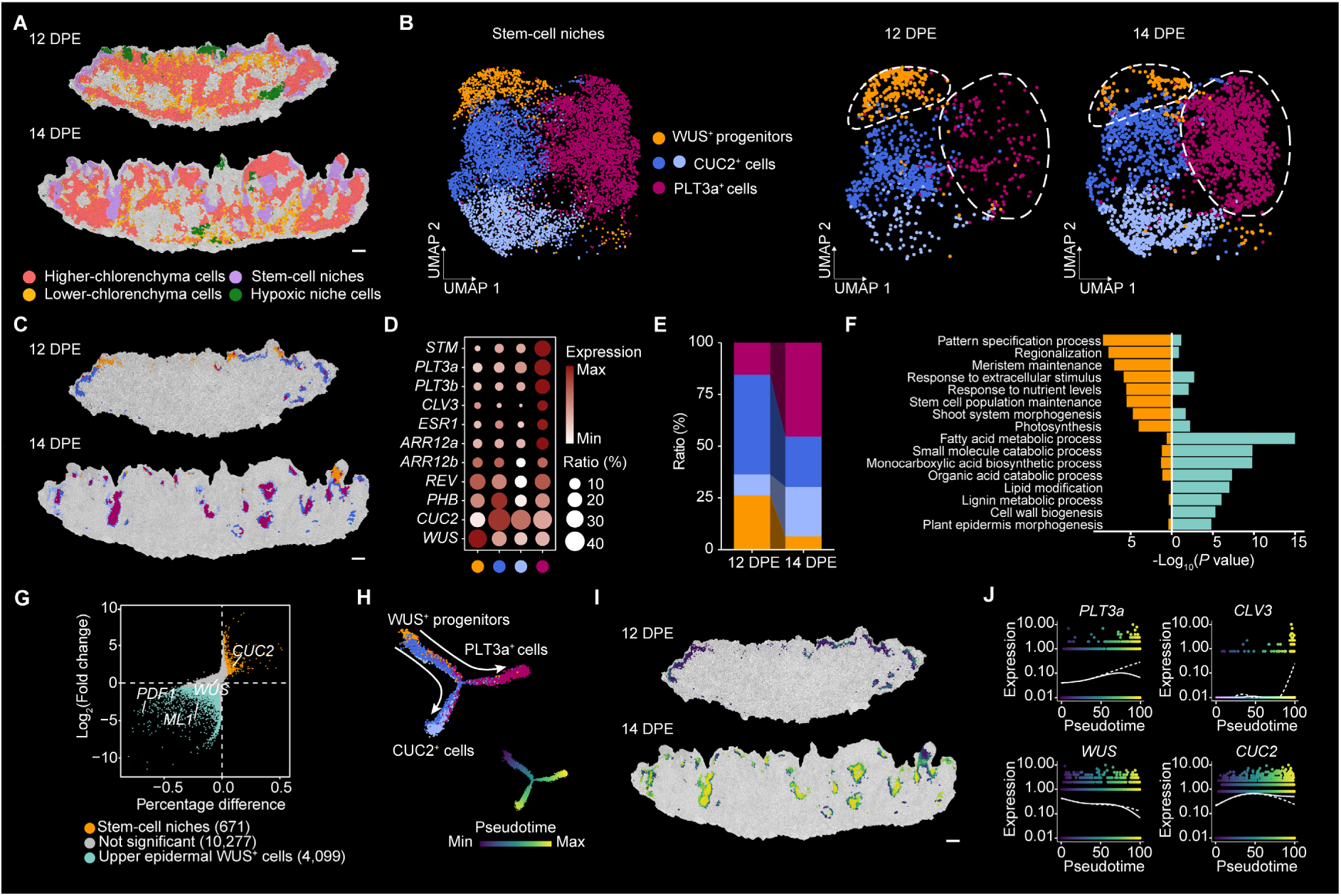
Sub-cell-types heterogeneity of tomato callus. **(A)** Spatial maps of the pluripotent cells at 12 and 14 DPE, showing higher- and lower-chlorenchyma cells, hypoxic niche cells, and SCN cells. Scale bar, 200 μm. **(B)** UMAP visualization of SCN sub-cell-types. Cells are colored by types, including WUS⁺ progenitors, CUC2⁺ peripheral cells, and PLT3a⁺ central cells. **(C)** Spatial distributions of niche sub-cell-types projected onto tissue sections at 12 and 14 DPE, colored as in **B**. Scale bar, 200 μm. **(D)** Bubble plot displaying the expression levels of marker genes for sub-cell-types in SCNs. **(E)** Proportions of niche sub-cell-types at 12 and 14 DPE, based on sequencing-based spatial transcriptomics (BMK S3000), colors as in **B**. **(F)** GO enrichment analysis of differentially expressed genes (DEGs), at least 2-fold changes, *P* < 0.01, between SCN WUS⁺ cells (orange) and upper epidermal WUS⁺ cells (cyan), based on sequencing-based spatial-transcriptomics (BMK S3000) spanning 8-14 DPE. **(G)** Scatter plot of DEGs used in **F**. Orange, upregulated in SCN WUS⁺ cells; cyan, upregulated in upper epidermal WUS⁺ cells; gray, not significant. Gene counts are annotated in brackets. **(H)** Pseudotime trajectory reconstruction of niche sub-cell-types, colored by identity (up) or pseudotime (down). **(I)** Spatial projection of pseudotime states onto callus sections at 12 and 14 DPE, colored as the pseudotime in **H**. Scale bar, 200 μm. **(J)** Expression trends of *PLT3a*, *CLV3*, *CUC2*, and *WUS* along pseudotime. Dashed/solid lines denote differentiation trajectories from WUS⁺ progenitors to PLT3a⁺ central cells and CUC2⁺ peripheral cells. Cells are colored by pseudotime in **H**.

Pseudo-time trajectory analyses [62] suggested a stem-cell differentiation hierarchy, showing that WUS^+^ cells diverge into inner and outer positional domains within the niche. The inner domain exhibits enhanced ribosome biogenesis and sequential marker activation, with *PLT3a* initiated early and *CLV3* activated later (namely PLT3a^+^ cells), whereas the outer domain is marked by late activation of *CUC2* (namely CUC2^+^ cells) (Figure 5B, 5H–5J, Supplemental Figure 6C). Time-course observations of *WUS* expression showed spatiotemporal co-expression with cyclins and cytokinin-related B-type *Arabidopsis Response Regulators* (*ARRs*) [69], potentially originating from cotyledon cambium-derived precursors (Supplemental Figure 4, 6D and 6F) [4]. Indeed, early regeneration stages show *WUS* activation in wound-adjacent regions and vascular tissues, later consolidating into niche foci (Supplemental Figure 4 and 6E) [70]. At early stages, *WUS* expression overlapped with *CUC2*, but at later stages *WUS* became restricted to the central domain while *CUC2* expression was enriched in the periphery (Figure 5J, Supplemental Figure 6E) [70]. Spatial co-expression networks placed *WUS* at the center of a module together with differentiation modulators *INCURVATA 4* (*ICU4*) and hormone-responsive genes *JASMONATE-ZIM-DOMAIN PROTEIN 1* (*JAZ1*) and *WOODEN LEG 1* (*WOL1*) (Supplemental Figure 6F). Our results show that early *WUS* expression in the upper epidermis, followed by its subsequent confinement to the central region of the regenerating SCN, is consistent with a spatial pattern shaped by regulatory constraints. However, such spatial shifts in gene expression may reflect either changes in the positions of the expressing cells or successive activation in neighboring cells, underscoring consistency with previous studies and highlighting the value of our dataset in providing super-resolution, whole-section, single-cell-level spatial data to disentangle these possibilities [70].

Taken together, these datasets define the vascular origin of callus, uncover pluripotent potential within upper-epidermal subtypes, and resolve the spatial stratification of WUS⁺ progenitors within the SCN, thereby capturing regeneration at high spatial and cellular resolution. The resulting multimodal single-cell spatial atlas serves as a standardized reference for benchmarking spatial transcriptomics and generating hypotheses on tissue origin and fate plasticity. As such, this resource provides both a methodological framework for addressing plant-specific challenges and a comprehensive reference for investigating regeneration across developmental contexts.

## DISCUSSION

Here, a spatial multi-omics approach generated a high-resolution atlas of tomato regeneration, enabling detailed characterization of cell-state transitions over space and time. The accompanying Plant Single Cell Spatial platform provides a user-friendly resource for further investigation into spatial gene regulation and niche dynamics in plant development. This study focuses on spatial omics and spatial architecture, and consequently does not deeply investigate auxin and cytokinin signaling, which have traditionally been the main focus of regeneration research [10, 17]. Our previous work has shown that these hormone signals tend to be broadly expressed [21], establishing a permissive regulatory background that operates in parallel to the spatially resolved mechanisms highlighted here.

The technical advances showcased in this study — integration of sequencing-based and imaging-based spatial transcriptomics — provide a resolution surpassing current single-cell techniques [19, 71]. Future applications include expanding this approach to other plant species and broader contexts, demonstrating the utility of this framework.

## ACKNOWLEDGMENTS

We acknowledge the support from the THU-IDG/McGovern Open Laboratory of Shared Instruments for Brain Science and the Cellular and Molecular Biology Platform, Beijing Research Institute of Chinese Medicine, Beijing University of Chinese Medicine. This work was supported by Shandong Provincial Natural Science Foundation (grant nos. SYS202206, ZR2023QC026 and ZR2023QC106), the National Natural Science Foundation of China (grant nos. 32200249 and 32170574), and the Taishan Scholars Program and Yuandu Scholars Program. No conflict of interest was declared.

## AUTHOR CONTRIBUTIONS

B.L., X.S., S.Z., Z.Y. conceived of the project and designed the experiments; S.Z., X.S., S.C., Y.N., M.L. and T.W. prepared the genetic materials; S.Z., and Z.Y. conducted the majority of experiments with contributions from S.C., Y.N., Y.S., L.X., Y.M., M.L., J.L., N.L., T.W., C.Q. and M.X.; X.S. and Y.L. performed the bioinformatics analysis with contribution from H.Y., Z.Y. and S.Z.; B.L., G.L., H.C., B.S., R.X., J.W., H.Z., S.H. and X.W.D. wrote the manuscript with input from all co-authors.

**Correspondence and requests for materials** should be addressed to B.L.

## METHODS

### Tomato materials and growth conditions

*Solanum lycopersicum* cv. Micro-Tom (MT, LA3911) was used as the wild-type background. Seeds were surface-sterilized by immersion in 75% (v/v) ethanol for 60 s, followed by shaking in 1% (v/v) sodium hypochlorite solution containing 0.02% (v/v) Triton X-100 for 15 min. The seeds were then rinsed eight times with sterile distilled water and dried on sterile filter paper. Sterilized seeds were germinated on half-strength Murashige and Skoog (1/2 MS) medium (2.16 g/L MS salts, 1% [w/v] sucrose, 3.5% [w/v] Phytagel, pH 5.8). Plates were incubated in the dark for 3 days, then transferred to long-day conditions (16 h light/8 h dark) at 25°C. Seedlings were grown in a growth chamber under long-day (LD) conditions at 25°C with 70% relative humidity and a light intensity of 50 µmol photons m⁻² s⁻¹.

### Shoot regeneration from tomato explants

To induce shoot regeneration, a 5-mm central segment was excised from the cotyledons of 6-day-old seedlings using a scalpel and placed adaxial side down in direct contact with shoot induction medium (SIM; 4.33 g/L MS basal salts, 3% [w/v] sucrose, 1 mg/L zeatin, 0.1 mg/L IAA, and 3.5% [w/v] Phytagel, pH 5.8). Cultures were maintained at 25°C under 70% relative humidity and a 16 h light/8 h dark photoperiod with a light intensity of ∼50 µmol photons m⁻² s⁻¹. Shoot regeneration efficiency was quantified as the average number of regenerated shoots per explant.

### Sequencing-based spatial-transcriptome methodology

#### Tissue cryosectioning

Fresh callus tissues were embedded in 100% pre-chilled OCT (Sakura, 4583) on ice. Infiltrated tissues underwent vacuum**-**assisted infiltration for 5 min at 4°C, followed by centrifugation at 3,000 × *g* (4°C, 5 min) to eliminate residual gas bubbles. The callus tissues were then transferred to 17 × 17 × 5 mm embedding molds (ShiTai, 80203-0007, China) containing pre-chilled OCT for embedding. The samples were adjusted, placed in a -80°C cryostat for OCT solidification, and stored at -80°C. The embedded frozen samples were sectioned at 10-μm thickness (Dakewe, CT520, China) for histological observation and spatial-transcriptome sequencing.

#### BMKMANU S1000 & S3000 RNA-seq

BMKMANU S1000 & S3000 RNA-seq were performed using the BMKMANU S1000 and BMKMANU S3000 gene-expression assay kits (BMKMANU, ST03002). Following the BMK Spatial Transcriptome User Guide (BMKMANU Gene Expression Kits Guide V3.2), procedures included tissue sectioning, Toluidine blue (Sigma, 6586-04-5) staining, 3D scanner image acquisition (3DHISTECH, Pannoramic MIDI II), tissue permeabilization (14 min), reverse transcription, cDNA first-strand denaturation, cDNA second-strand synthesis, cDNA second-strand denaturation and recovery, cDNA amplification purification, and library construction. Illumina libraries were sequenced using Illumina NovaSeq.

#### BGI Single-Cell Stereo-seq

The Stereo-seq process followed the Stereo-seq Transcriptome Assay Kit (Carrier Version) User Guide. Tissue sections were first adhered to the Stereo-seq chip T-carrier surface, incubated at 37°C for 1 min, and then fixed in pre-chilled methanol at -20°C for 1 h. Subsequently, tissue sections were stained for cell walls with Fluorescent Brightener 28 (FB) and for cell nuclei with Qubit ssDNA HS (ssDNA, Invitrogen, Q10212). Imaging was conducted using an automated fluorescence chip scanning system (Leica, DM6B). Procedures included tissue permeabilization, reverse transcription, tissue removal, cDNA release and recovery, cDNA release and amplification, and library construction. Spatial libraries were sequenced on the DNBSEQ-T10 sequencer.

### SmFISH using ISSeq-based spatial transcriptomic Kit

Fresh tissues were sectioned into 10-µm slices and then mounted onto pretreated glass-bottom plates. First, the glass slide was transferred from dry ice to a 37°C pre-heated thermal cycler and held for 1 min. Next, the slices were fixed with 4% v/v paraformaldehyde (PFA) in phosphate-buffered saline (PBS) at room temperature for 10 min. After that, slices were rinsed three times with 1x PBS for 1 minute each and permeabilized with 0.3% v/v Triton X-100 for 30 min prior to hybridization. Subsequently, the slices were gently washed with PBSTR for 1 min, and this washing step was repeated three times. Padlock probes were dissolved at a concentration of 50 µM in ultrapure RNase-free water. Firstly, the probe mixture was heated to 95°C for 5 min and then immediately cooled on ice for 2 min. Secondly, the probe-hybridization mix was prepared, which consisted of a final probe concentration of 1 µM along with RCA primer (at a 1:1 molar ratio, from the ISSeq Kit v1.0, BMKMANU, China), 10% v/v formamide, 4× saline sodium citrate (SSC), and 2 U RNase inhibitor (from the ISSeq Kit v1.0, BMKMANU, China). Thirdly, the probe-hybridization mix was carefully added along the side of the well to ensure uniform coverage of the tissue sections without the introduction of bubbles, followed by incubation at 45°C for 4 h. After incubation, all probe-hybridization mix was removed, and PBSTR was added. The slices were incubated at room temperature for 1 min and then washed with hybridization wash buffer (4x SSC, 1x PBSTR) at 37°C for 20 min each. Finally, the sample was briefly rinsed twice with PBSTR at room temperature. Subsequently, the samples were incubated with ligation mixture (containing 75 U ligase, 1× ligase buffer, and 2 U RNase inhibitor (from the ISSeq Kit v1.0, BMKMANU, China)) at room temperature for 2 h. After ligation, the sample was briefly rinsed twice with PBSTR at room temperature. Then, the samples were incubated with RCA mixture (containing 75 U DNA polymerase, 500 µM dNTP, 1× DNA polymerase buffer, and 1 U RNase inhibitor (from the ISSeq Kit v1.0, BMKMANU, China) at 30°C for 12 h. Once again, the sample was briefly rinsed twice with PBSTR at room temperature. Finally, fluorescence imaging using a microscope (BH1000, BMKMANU, China), the fluorescent oligonucleotide complementary to the DNA amplicon was diluted to 500 nM in hybridization mix (4x SSC, 40% v/v formamide) (from the ISSeq Kit v1.0, BMKMANU, China). The samples were incubated at room temperature for 10 min and then washed three times with 1x PBS prior to imaging. Before imaging, a coverslip was placed over the sample, and the sample was stained with DAPI.

### Xenium *in situ* gene expression

#### Formalin-fixed paraffin embedding

After trimming the experimental samples to appropriate dimensions, immerse them in pre-cooled FAA fixative for initial fixation. Perform vacuum infiltration for twice. Replace with fresh fixative and incubate at 4°C for 12 h for thorough fixation. Dehydrate the samples using a graded ethanol series (50%–100%, v/v; 30 min per step). Following dehydration, perform ethanol-xylene gradient substitution: sequentially transfer through 30%, 50%, 70%, 100% xylene I, 100% xylene II, and 100% xylene III (30 min per step). Pre-infiltrate the samples with 50% v/v paraffin to ensure complete dissolution of the paraffin. Gradually supplement the paraffin and replace with fresh solution every 3 h for a total of three replacements. Embed the samples, and store the paraffin blocks at 4°C for subsequent use.

#### Preparation of sequential sections

Sequential sections were prepared according to manufacturer instructions (‘Tissue Preparation Guide Demonstrated Protocol CG000578’ for Xenium). Cut the tissue block at 15-µm until all of the edges of the tissue are exposed or until the region of interest is exposed. Five-micrometer sequential sections were then collected, floated in a 37_°_C water bath, and adhered to Xenium slides (10x, PN 1000460). Calli were sliced as close to the center of the active area as possible for Xenium slides. Samples were baked at 42_°_C for 3 h and sections were stored in a desiccator at room temperature according to the manufacturer’s instructions prior to processing.

#### 10x Xenium data acquisition

10x Xenium samples were processed in three batches according to manufacturer protocols “Probe Hybridization, Ligation & Amplification, User Guide CG0000582” and “Decoding & Imaging, User Guide CG000584”. The Xenium custom gene-expression panel containing probes for 50 tomato genes were applied on each section to perform hybridization, ligation and rolling-circle amplification, followed by cycles of fluorescent-probe hybridization, imaging and decoding. Slides were processed on a 10x Xenium Analyzer, with ROIs (Region of interest) selected to cover the entire callus region. Data were processed using the standard 10x workflow to generate cell-by-gene and transcript-by-location matrices. The data were visualized by Xenium Explorer v3.2.

### Spatial or single-nucleus RNA-seq data analysis

#### BMKMANU S1000/S3000 RNA-seq

Raw data were mapped to the tomato reference genome (version SL4.0 and Annotation ITAG4.1) using BSTMatrix (v2.4) to obtain gene-expression count matrices of different resolutions. The histological images obtained from spatial experiments were used for cell segmentation through automatic recognition and manual correction using BSTCellViewer (v4.1.12) software. The gene-expression count matrix with single-cell resolution obtained from BSTCellViewer software was processed using the R package Seurat (v5.1.0) [72].

#### BGI single-cell Stereo-seq

The ScStereo-seq processes were performed according to a previously reported method [72] with minor modifications. Raw data were mapped to the tomato reference genome using SAW (v8.0) to obtain gene-expression count matrices of different resolutions. Histological images obtained from spatial experiments were used for cell-segmentation through automatic recognition and manual correction using BSTCellViewer (v4.1.12) software. The gene-expression-count matrix with single-cell resolution obtained from SAW software was processed using the R package Seurat (v5.1.0).

#### Single-nucleus RNA-seq

Raw data from 10x Genomics and BMKMANU DG1000 were aligned to the tomato reference genome using CellRanger (v7.0.0) and BSCMatrix (v2.1), respectively. The obtained gene-expression-count matrix was subsequently clustered and annotated using R package Seurat (v5.1.0) [72].

#### Clustering of spatial or single-nucleus RNA-seq data

To remove low-quality data, we filtered cells with gene numbers > 30 and genes that were expressed in > 10 cells. Seurat (v5.1.0) [72] was used to normalize, scale, and transform data. Harmony (v1.2.0) [73] with the default parameters was used to correct batch effects. For dimension reduction of the data, principal-component analysis (PCA) was conducted and the top-30 principal component were retained for further analysis. Cell clustering based on Louvain (“FindNeighbors” and “FindClusters”) and visualization with non-linear dimensional-reduction algorithms (“RunUMAP”) were performed. The differentially expressed genes (DEGs) between clusters were calculated using “FindAllMarkers” or “FindMarkers” functions. The following cutoffs were applied: only.pos = TRUE, logfc.threshold = 0.5, min.pct = 0.1). Spatial visualization of cell clusters and gene expression on cell masks were performed by customized Python scripts for spatial-transcriptome data (https://github.com/Tako-liu/lilab_spatial/tree/main).

#### Joint clustering spatial and single-nucleus RNA-seq datasets

To provide one consensus cell-type taxonomy based on both spatial and single-nucleus RNA-seq datasets of more than 1 million cells, we optimized the clustering pipeline to integrate datasets collected by different transcriptome-analysis platforms, thereby aiming to build a graph that incorporates samples from all the datasets. Spatial RNA-seq data were selected as the reference dataset, and the clustering pipeline above was used to build an integrated reference. Subsequently, this integrated reference was employed to annotate datasets from other platforms. The pipeline was executed through two steps: Firstly, the selection of anchor cells for both the reference and query datasets was achieved by employing the FindTransferAnchors function. Secondly, upon the identification of these anchor cells, the TransferData function was then applied to classify the query cells based on the characteristics derived from the reference dataset.

#### Pseudo-time analysis

Monocle2 (v2.30.1) [74] and Monocle3(v1.3.7) [75] were used to generate a pseudo-time trajectory and identify developmentally variable genes using cluster information previously identified using Seurat. Briefly, the raw Seurat object was first converted into a monocle2 or monocle3 object. The DEGs between clusters were used for ordering genes in monocle2. Then, we performed dimensionality reduction analysis on the data using the “DRTree” method and performed trajectory-inference analysis using default parameters in monocle2. Developmentally variable genes were identify using “differentialGeneTest” function. The ‘UMAP’ method was used for data dimensionality reduction analysis in monocle3.

#### RNA-velocity analysis

RNA velocity of spatial RNA-seq data was evaluated by velocyto package [62] and Dynamo (v1.4.0) [74]. The loom file containing splicing information was generated with default parameters by velocyto package. The rates of gene splicing and degradation were estimated by analyzing the relative abundance of nascent (unspliced) and mature (spliced) mRNA using the Dynamo package. The unspliced and spliced raw count matrices of specific cell types were extracted and processed using the cell_monocle function. The UMAP, spatial coordinates, and cluster information extracted by Seurat were also embedded in Dynamo. Then, kinetic parameters and gene-wise RNA velocity vectors were estimated on the normalized matrix using the cell_velocities function and then projected into a visualized UMAP.

#### Gene co-expression network analysis

To identify gene-expression relationships, hdWGCNA [76] was employed for co-expression network analysis. Default parameters of hdWGCNA were applied to the analysis of the integrated gene-expression-count matrix. Briefly, genes with detectable expression in at least 2% of cells were incorporated into the network construction, ensuring that all cells within the matrix were considered. For the construction of metacells, Harmony [73] was applied for cell-dimension reduction and cell-type annotation. The parameters were set to a minimum of 30 cells and a maximum of 10 shared genes per metacell. From each module, the top-30 genes exhibiting the highest TOM (topological overlap matrix) values associated with the target gene were identified and used in network construction. This strategy was employed to delineate the regulatory network interactions between the target gene and other genes within the module.

#### Gene-ontology (GO) enrichment analysis

Based on the known gene function and GO biological process in *Arabidopsis*, the homologs of *Arabidopsis* genes (TAIR10) in the tomato genome were identified using BLASTP. GO enrichment analysis was performed using the R package clusterProfiler [77] with TAIR10 annotation as the background.

#### Distance analysis

To quantify the spatial distance between the queried cell type and subepidermal cells, a file containing the central coordinates of each cell was created for every sample using customized Python scripts (https://github.com/Tako-liu/lilab_spatial/tree/main). The files were read using R. Then, the distance for each queried cell to the subepidermal cells was ascertained by averaging the Euclidean distances to its three closest neighbors. The median of these distances was then adopted to represent the distance between the two distinct cell types. Visualization of the distance between the two cell types was performed using ggplot2.

### Bulk mRNA-seq data analysis

The mRNA sequencing (mRNA-seq) data analysis was performed as described in our previous study [78]. Briefly, total RNA was extracted using the RNeasy Plant Mini Kit (Qiagen). Libraries were prepared using an mRNA preparation kit, following the standard protocol. Sequencing of paired-end 150-base-pair mRNAs was conducted using MGI SEQ-2000 platform. Raw reads were filtered by fastp (v0.22.0) [79] with default parameters. Filtered reads were then aligned to the tomato genome (version SL4.0 and annotation ITAG4.1) using the hisat2 (v2.2.1) [80] program with two mismatches allowed. The differentially expressed genes were identified using the DESeq2 package in R [81].

Spatial transcriptomics data were generated at the Single-Cell and Single-Molecule Core Facility, Peking University Institute of Advanced Agricultural Sciences, using the 10x Genomics Xenium, BMKMANU S1000/S3000, and BGI Stereo-seq platforms.

---

**Supplemental Figure 1.**
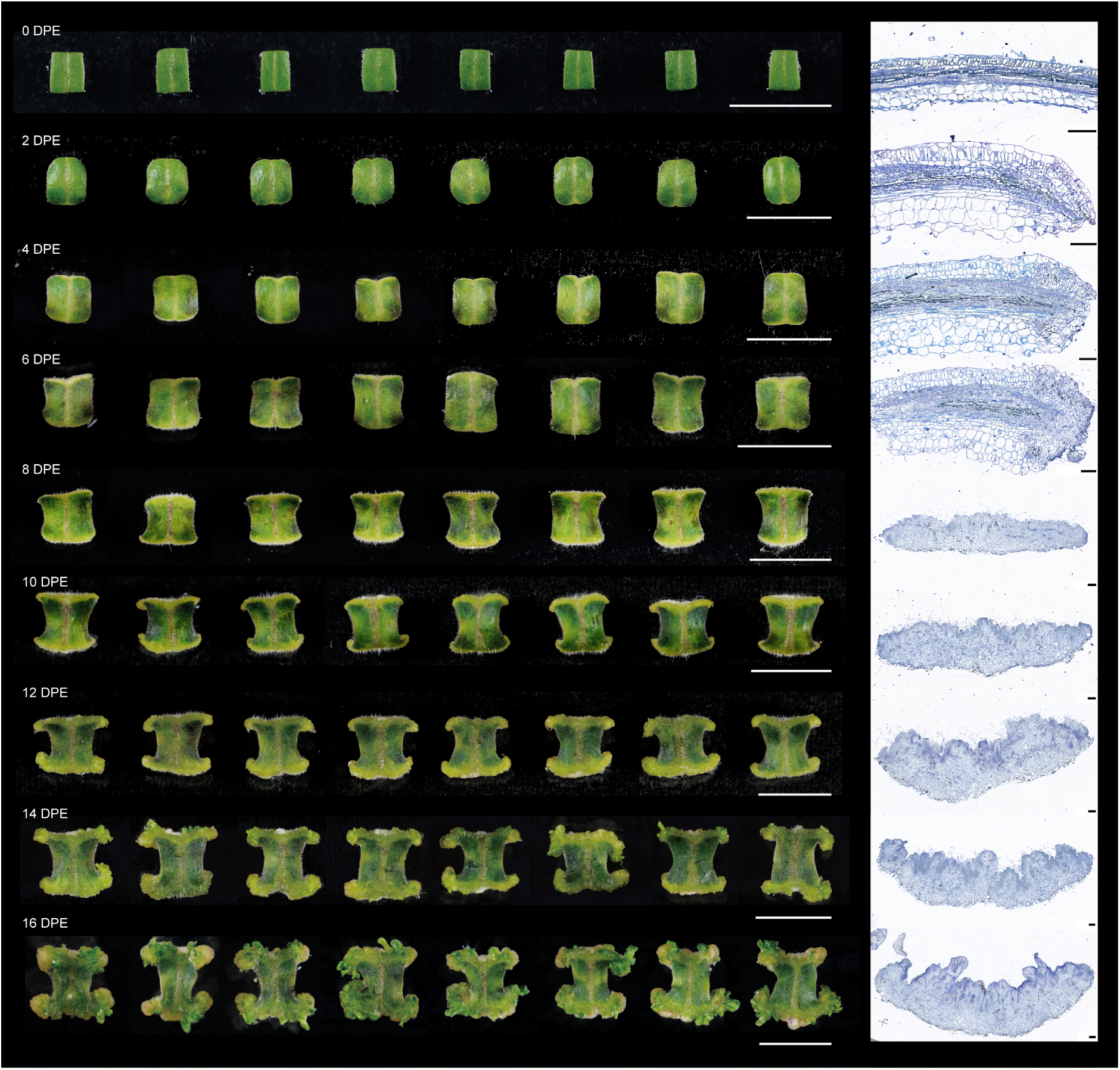
A well-established *de novo* regeneration system of tomato cotyledon explants. Representative images display the stages of tomato callus development. Photo images, scale bar, 1 cm. Sections with TBO staining, scale bar, 200 mm. Sectioning planes are indicated with dashed white lines in Figure 1A.

**Supplemental Figure 2.**
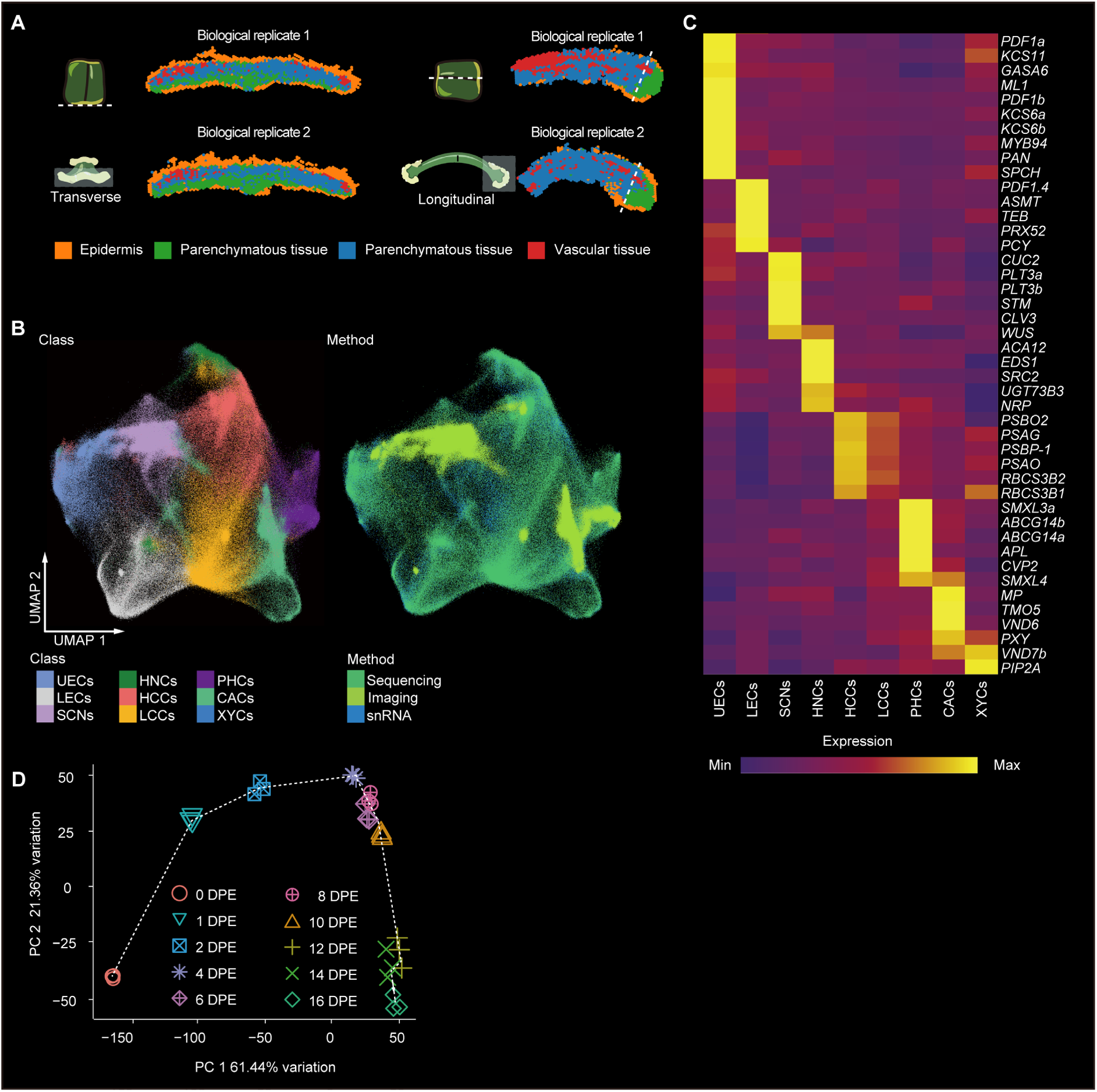
Integrating multimodal and spatiotemporal transcriptomics of tomato regeneration. **(A)** Schematics for transverse and longitudinal cutting of 6 DPE callus, as well as demonstrating consistency between replicates. **(B)** Integrated uniform manifold approximation and projection (UMAP) visualization of all profiled cells obtained from sampling as in (Figure 1A). Left: Cell-type classification: upper epidermal cells (UECs), lower epidermal cells (LECs), stem-cell niches (SCNs), hypoxic niche cells (HNCs), higher-chlorenchyma cells (HCCs), lower-chlorenchyma cells (LCCs), phloem cells (PHCs), cambium cells (CACs), and xylem cells (XYCs). Right: integrated datasets: sequencing-based spatial transcriptomics (Sequencing), imaging-based spatial transcriptomics (Imaging) and snRNA-seq (snRNA). **(C)** Heatmap of sequencing-based spatial expression levels of known marker genes. Cell-type abbreviations are as in the **B**. **(D)** Principal-component analysis (PCA) of bulk RNA-seq data (three replicates per time point). Shapes denote sampling time points; dotted arrows indicate progression trajectory.

**Supplemental Figure 3.**
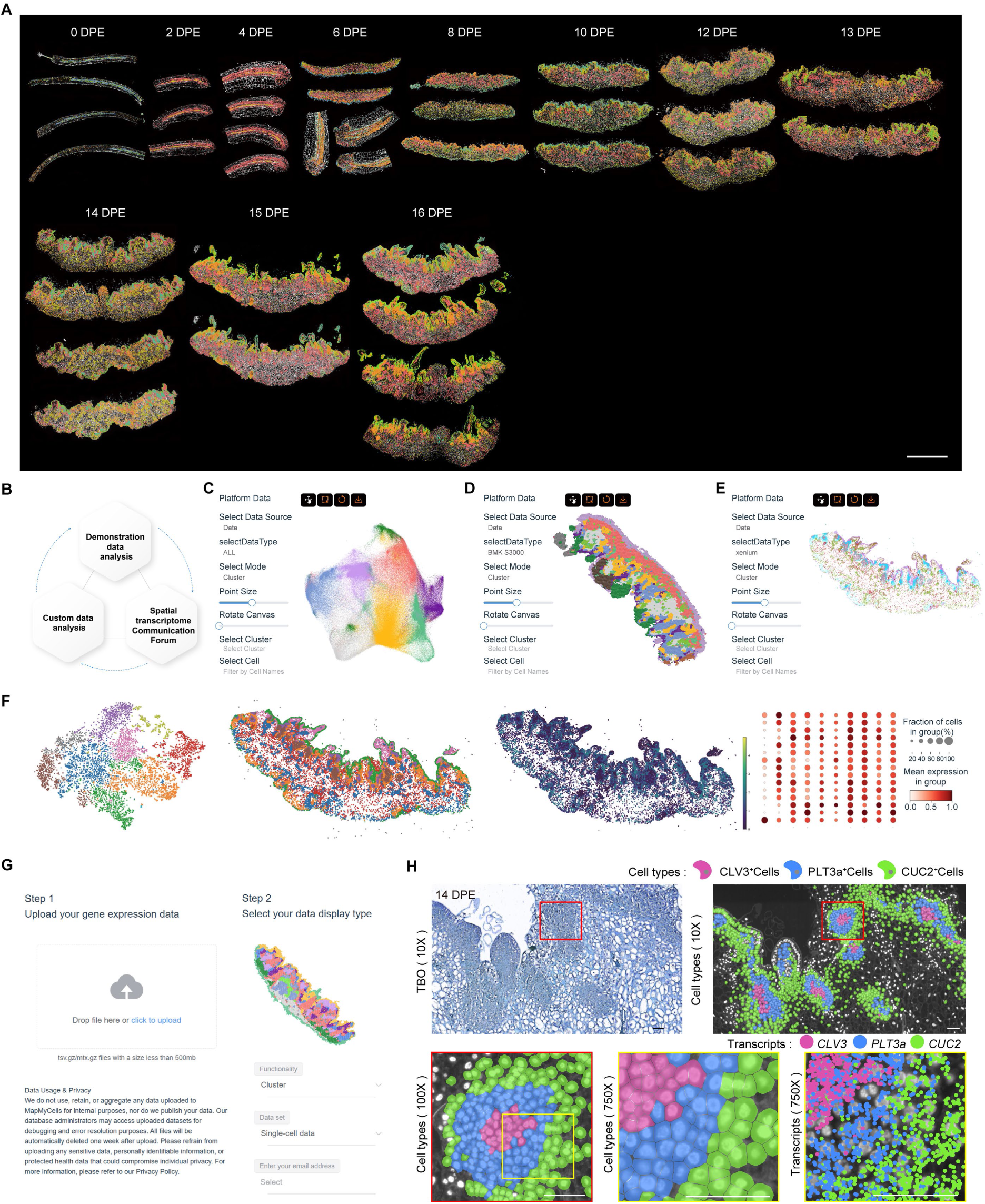
Reproducibility assessment of the imaging-based spatial transcriptomics and overview of the Plant Single Cell Spatial Platform. **(A)** Paraffin embedding maintains tissue integrity, ensuring high reproducibility between replicates in imaging-based spatial transcriptomics. Scale bar, 2 mm. **(B**-**G)** Plant Single Cell Spatial Platform. (**B**) Overview of the Plant Single Cell Spatial Platform’s features. (**C**) Screenshots of UMAP clustering visualization. (**D**) Spatial clustering visualization. (**E**) Clustering visualization of imaging-based data. (**F**) Output examples of the online spatial-transcriptomics analysis tool. (**G**) Data-upload interface for customized analysis. **(H)** Subcellular-resolution profiling of tomato SCN structure using 10x Xenium at increasing magnifications. Markers: *CLV3* (core, cells in magenta, transcripts in pink), *PLT3a* (inner, cells in blue, transcripts in blue), *CUC2* (shell, cells in green, transcripts in green). Scale bar, 50 μm.

**Supplemental Figure 4.**
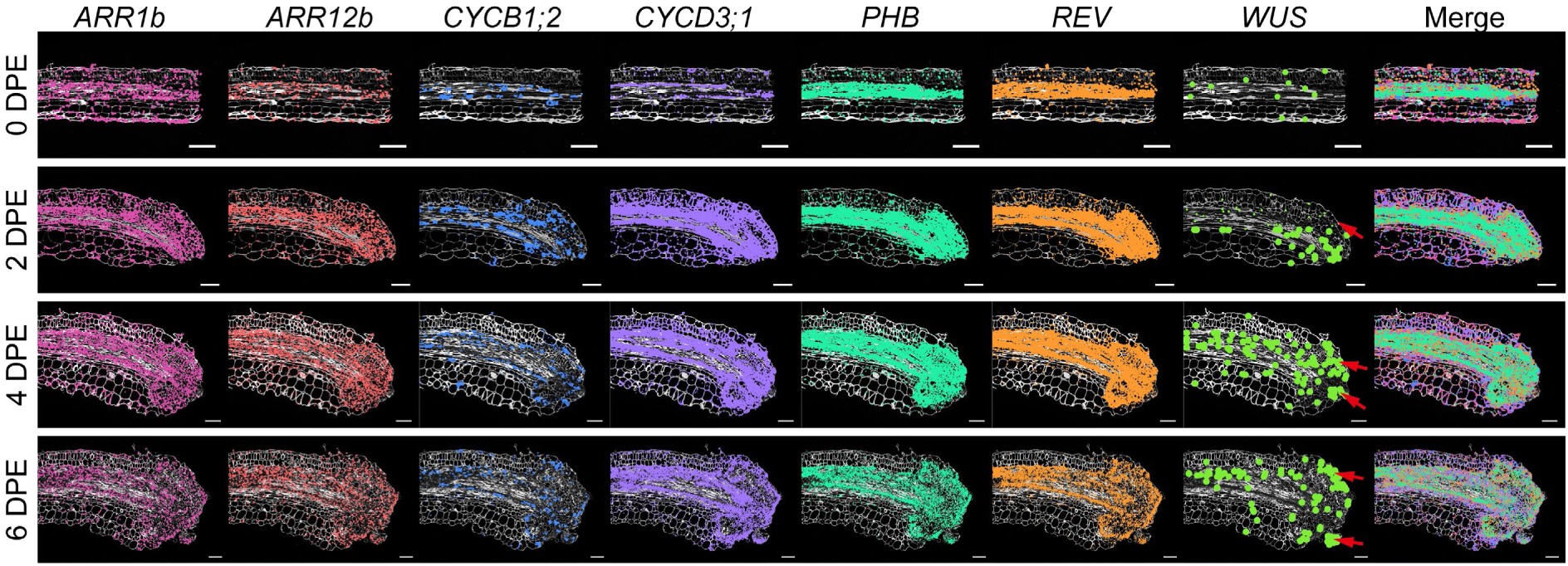
Marker gene expression during callus initiation. 10x Xenium spatial plots showing the transcripts of *ARR1b* (deep pink), *ARR12b* (light red), *CYCB1;2* (blue), *CYCD3;1* (purple), *PHB* (light green), *REV* (orange), and *WUS* (green, highlighted by larger dots) transcripts during the early stages (0-6 DPE) of regeneration. Scale bar, 100 μm.

**Supplemental Figure 5.**
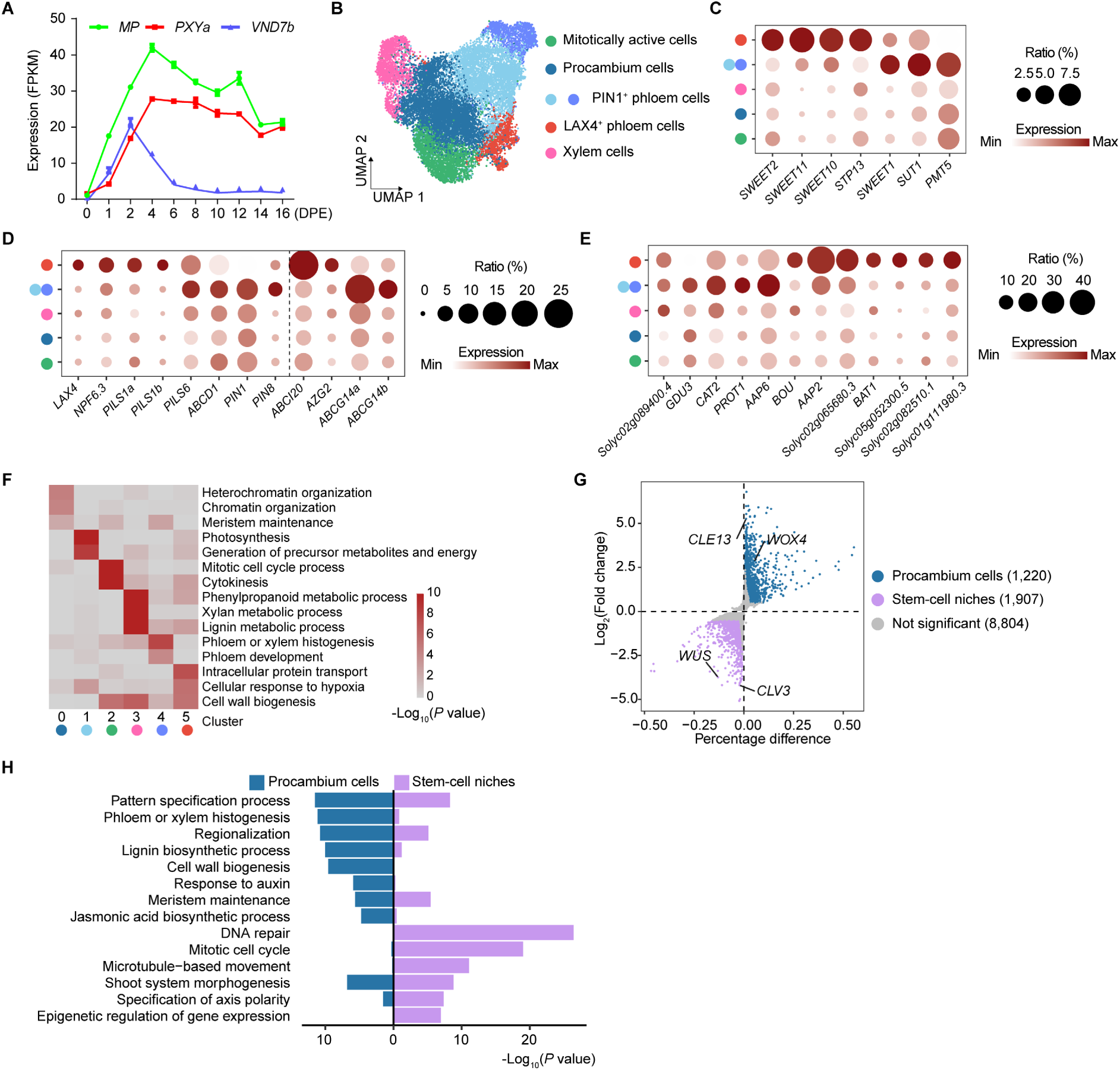
Characterization of vascular cell sub-types during tomato callus regeneration. **(A)** Bulk RNA-seq profile of expression for vascular regulators (*MP*, *PXYa*, and *VND7b*) during regeneration (0-16 DPE). Error bars indicate mean ± SD (n = 3 biological replicates). **(B)** UMAP visualization of sequencing-based spatial transcriptomics, identifying five vascular sub-cell-types: procambium cells, mitotically active cells, xylem cells, PIN1⁺ phloem cells, and LAX4⁺ phloem cells. **(C-E)** Bubble plots showing the expression levels of genes associated with glucose transport **(C)** phytohormone transport (**D**) and amino acid transport (**E**) across vascular tissue sub-cell-types. The colors of the point on the left are consistent with **B**. **(F)** GO enrichment analysis of genes specific to vascular-tissue cells. The colors of the point on the bottom are consistent with **B**. **(G)** DEGs between cell clusters of procambium and SCN from the integrated sequencing-based spatial transcriptomics of 8-14 DPE. Purple dots represent genes with high expression in SCNs, while steel blue dots denote genes highly expressed in procambium cells. **(H)** GO enrichment analysis of DEGs in **G**, showing significant pathways such as pattern specification, lignin biosynthesis, auxin response, and shoot system morphogenesis.

**Supplemental Figure 6.**
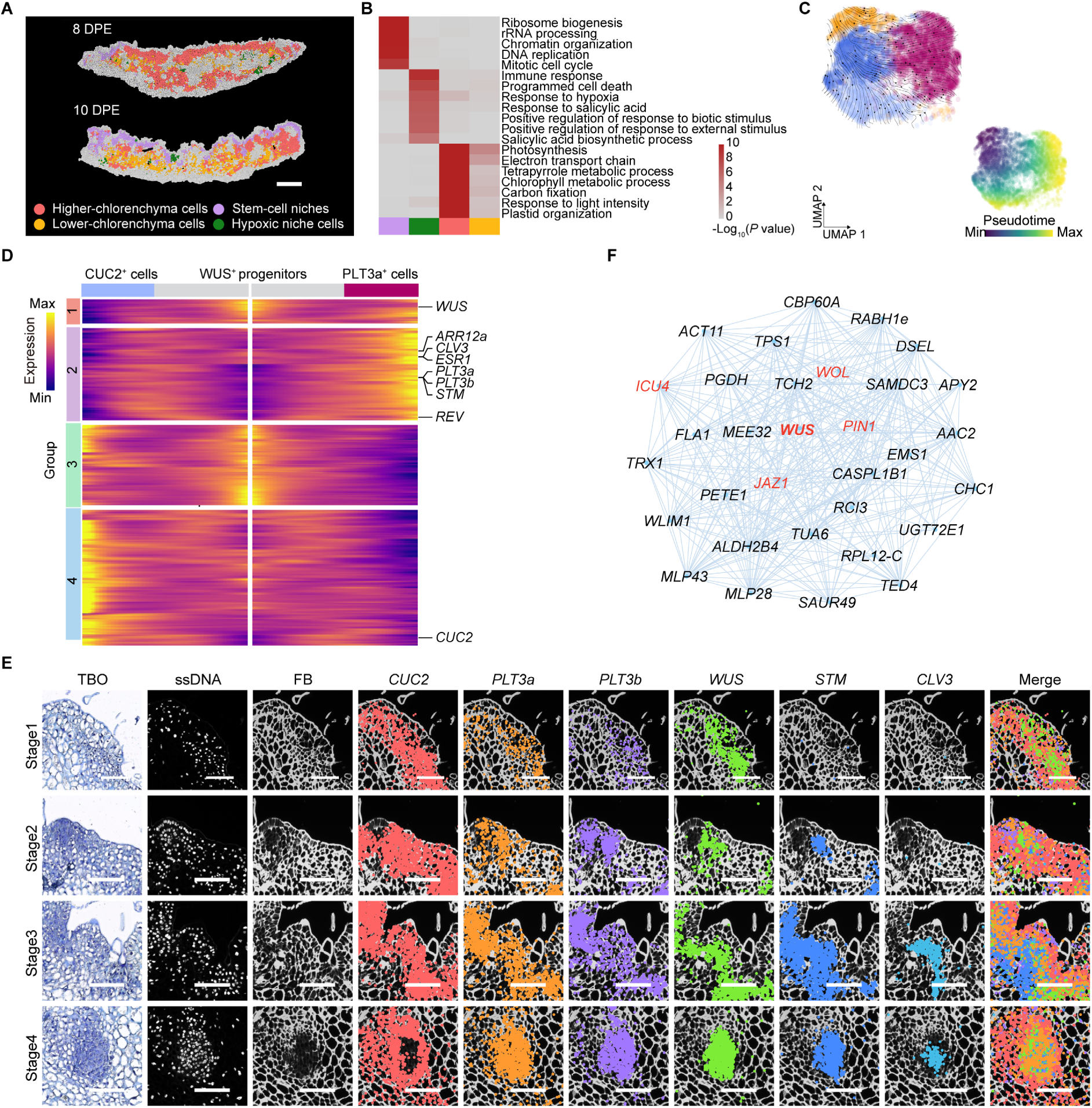
SCN formation and functional differentiation. **(A)** Spatial maps of pluripotent cells at 8 and 10 DPE, including stem-cell niches, hypoxic niche cells, higher-chlorenchyma cells, and lower-chlorenchyma cells. Scale bar, 500 μm. **(B)** GO enrichment analysis of genes associated with pluripotent cell types in **A**. **(C)** RNA velocity and pseudotime trajectory reconstruction of SCN sub-cell-types, colored by identity (up) or pseudotime (down). **(D)** Heatmap of gene expression along pseudotime of SCN cells, grouped (group 1-4) by their expression dynamics during lineage progression. Known stemness regulatory genes are annotated at right. **(E)** 10x Xenium spatial plot mapping transcripts of *CUC2* (light red, highlighted by larger dots), *PLT3a* (orange), *PLT3b* (purple), *WUS* (green), *STM* (blue), and *CLV3* (sky blue) in SCNs at different developmental stages. Scale bar, 30 μm. **(F)** Co-expression network centered on *WUS*, showing links to differentiation regulators (*ICU4*, *JAZ1*, *WOL*) and hormone-responsive gene (*PIN1*)

